# Synthesis of Plasma Membrane-Enriched Nanodiscs from Giant Plasma Membrane Vesicles

**DOI:** 10.64898/2026.11.20.768238

**Authors:** Ciara O. Jacobs, Ryan R.S. Santee, Brandon S. Pizarro Carbajal, Wade F. Zeno

## Abstract

Nanodiscs are highly versatile biomaterials with applications in cell membrane mimicry, membrane protein (MP) characterization, and hybrid biological-abiological material synthesis. The plasma membrane (PM) is an ideal starting material for nanodisc synthesis because it naturally houses a diverse array of functionally important MPs in their native lipid environment. However, the PM comprises a small fraction of the total membrane material in a typical eukaryotic cell. Traditional methods to isolate the PM suffer from undefined relative PM enrichment compared to other cellular membranous structures. Here, we introduce a method to synthesize PM-enriched nanodiscs (PMNDs) from giant plasma membrane vesicles (GPMVs). We show that GPMV-derived PMNDs retain functional membrane-anchored protein, exhibit uniform size and morphology, and bind specifically and dose-dependently to the PM of cells expressing a cognate receptor. This strategy offers broad applicability for synthesizing nanodiscs with novel MPs for investigating PM-nanomaterial interactions, cellular targeting, and vaccine development using immunologic MPs.

## INTRODUCTION

Membrane proteins are involved in many dynamic cellular processes, including signal transduction^1^, enzymatic reactions^2^, and molecular transport.^3^ However, their innate reliance on native lipid environments previously made membrane proteins difficult to study and employ for therapeutic applications.^4,5^ In the past few decades, nanodiscs – discoidal patches of lipid bilayers stabilized by amphipathic scaffolds – have emerged as a powerful tool for solubilizing membrane proteins^6^ for structural characterization^7,8^, protein-lipid interaction studies^9–11^, and targeted therapeutic delivery.^12^ Traditional nanodiscs are synthesized using a wide range of synthetic lipids^13,14^ and are most often stabilized by helical membrane scaffold proteins (MSPs), such as MSP1D1, a truncation mutant of the human apolipoprotein A-I.^15^ Recently, numerous research groups have also successfully synthesized nanodiscs from native cell membranes primarily using polymer scaffolds. These polymer-stabilized cell-derived nanodiscs demonstrate broad applicability across high throughput organellar proteomics^16^, cancer immunotherapy^17^, and platelet-mimicking nanomedicine.^18^ One commonly used polymer scaffold is styrene maleic acid (SMA), an amphiphilic thermoplastic widely used in industrial coatings which has been shown to efficiently solubilize membrane proteins from cell membranes.^19^

Traditionally, cell-derived nanodiscs are synthesized from whole cell lysate which can be further purified to isolate specific organellar membranes.^16,20^ Common methods for isolating the plasma membrane from in-tact whole cells involve lysis by freeze-thaw cycles or homogenization. The harsh conditions imposed by these methods can lead to the delicate membrane proteins denaturing due to excessive shear forces or high temperatures.^21^ These methods of whole-cell mechanical disruption release both plasma and intracellular membranes, necessitating subsequent fractionation to enrich the plasma membrane.^22,23^ Therefore, conventional plasma membrane isolation methods suffer due to contamination from internal organelles and other intracellular structures. With the exception of red blood cells which are devoid of organelles, the plasma membrane represents less than 5% of a eukaryotic cell’s total membrane material, making it difficult to isolate with high purity.^5^ To isolate the plasma membrane from other membranous structures, differential or density gradient ultracentrifugation are often used^24–27^, but these methods suffer from poor scalability and the exact enrichment of the plasma membrane relative to other membranous structures in the whole cell homogenate remains unknown. Due to these shortcomings in cell lysis and plasma membrane enrichment, we propose giant plasma membrane vesicles (GPMVs) as an advantageous starting material for generating cell-derived nanodiscs, as they provide a plasma membrane-enriched fraction with minimal contamination from internal organelles.

GPMVs are spherical, micron-sized vesicles which can be isolated from live cells using osmotic shock or chemical vesiculation. They conserve the native protein-lipid composition of the plasma membrane, which is critical for maintaining the function of delicate membrane proteins, making them ideal isolating the plasma membrane for a wide range of applications. Additionally, previously developed protocols efficiently generate highly pure samples of GPMVs without contamination from other membranous structures or cytoskeletal fragments as observed by electron microscopy (EM).^28–30^ As a result, GPMVs are an ideal, novel starting material for nanodisc synthesis. Here, we present a platform to synthesize “Plasma Membrane-enriched Nanodiscs (PMNDs)” derived from GPMVs to ensure that nanodiscs are synthesized from a plasma membrane-enriched starting material.

To generate GPMVs, we treated adherent HeLa cells with N-Ethylmaleimide, a commonly used vesiculation agent^29^, to induce plasma membrane blebbing and vesicle formation. After sufficient incubation, GPMVs were isolated from their parent cells and further purified. Purified GPMVs were then processed and incubated with scaffolding proteins or polymers to yield PMNDs. By harvesting GPMVs directly from adherent cells, this approach avoids whole-cell mechanical disruption prior to membrane isolation. The resulting nanodiscs are derived from a plasma membrane-enriched starting material, retaining a complex cell-derived lipid environment that can support membrane protein stability and downstream applications (Figure 1).

**Figure 1.**
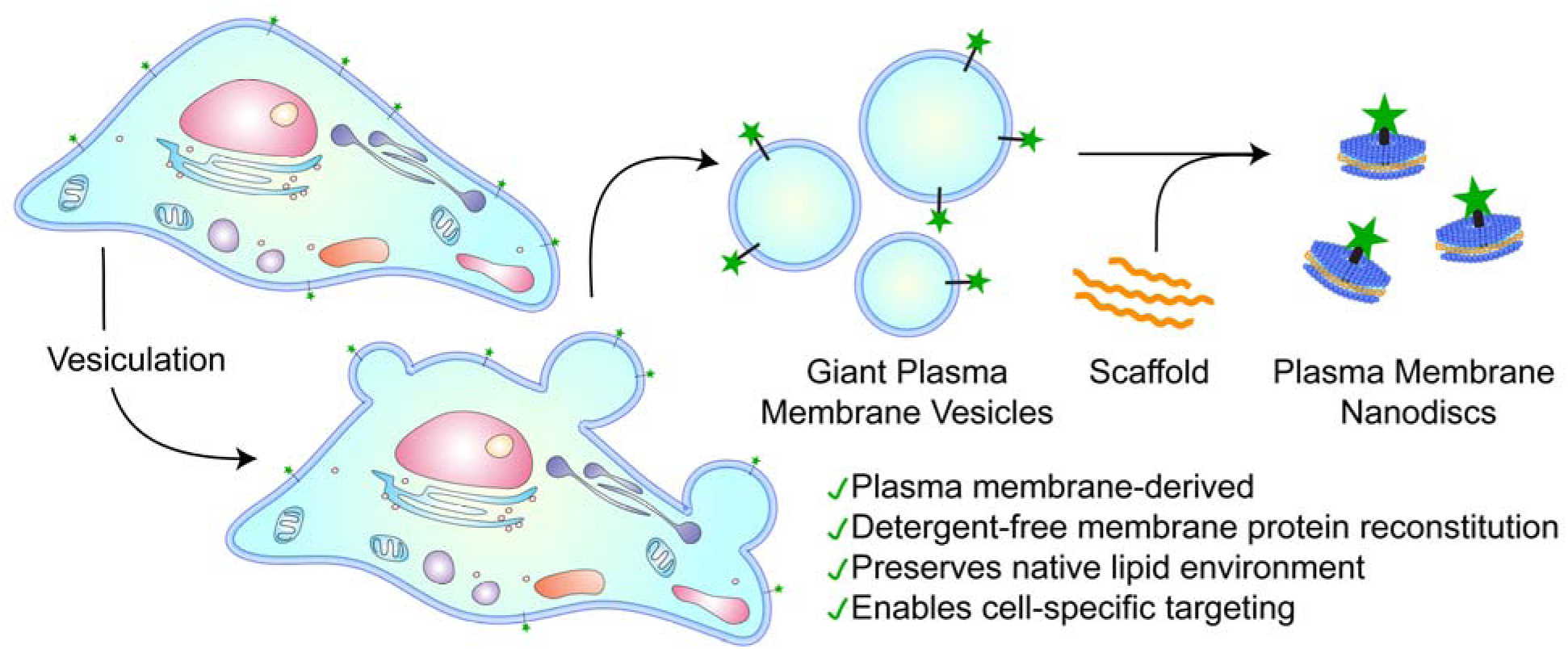
Schematic of PMND synthesis process. HeLa cells are incubated with N-ethylmaleimide to induce formation of giant plasma membrane vesicles (GPMVs). GPMVs are purified, processed, and incubated with amphipathic scaffold molecules to form plasma membrane-enriched nanodiscs (PMNDs).

## RESULTS AND DISCUSSION

We first aimed to optimize a detergent-free synthesis process for PMNDs by using different scaffolding species and varying membrane curvature of the starting vesicle populations. To modulate the curvature of the starting plasma membrane-derived vesicle populations, we briefly subjected GPMVs to probe sonication for 1 minute to generate sonicated plasma membrane vesicles (SPMVs). The two scaffolding species investigated were SMA and MSP1D1, hereafter referred to as MSP. Using these two scaffolds and two vesicle preparations differing in membrane curvature, we generated four PMND synthesis variants: (1) GPMVs + SMA, (2) GPMVs + MSP, (3) SPMVs + SMA, and (4) SPMVs + MSP (Figure 2A). We first sought to determine a metric for approximating membrane content and tuning the scaffold-to-membrane ratio. We determined the concentration of GPMVs (i.e., number of vesicles per unit volume) by counting vesicles within a defined volume using a hemocytometer (Figure 2B) and measured the average GPMV diameter from brightfield microscopy images using ImageJ (Figure 2C). We then used the estimated total GPMV surface area to calculate the scaffold concentration used for incubation, corresponding to approximately 1 scaffold molecule per 5 nm^2^ of membrane surface area. After overnight incubation and subsequent purification, PMND formulations were characterized using dynamic light scattering (DLS). Number-weighted size distributions demonstrated a marked size decrease in PMND formulations originating from both GPMVs and SPMVs (Figure 2D). However, both formulations derived from GPMVs exhibited multimodal size distributions, indicating high degrees of sample heterogeneity. PMNDs derived from SPMVs + MSP also exhibited heterogeneity, while PMNDs derived from SPMVs + SMA exhibited a more unimodal size distribution, having a strong peak near the expected hydrodynamic diameter value of 20 nm. PMND size, morphology, and homogeneity were further evaluated using negative stain transmission electron microscopy (TEM). TEM micrographs of both GPMV-derived formulations and SPMVs + MSP PMNDs revealed nanodisc structures alongside substantial amounts of non-nanodisc material (Figure 2E), likely consisting of residual scaffold and/or lipid-associated debris. PMNDs derived from SPMVs + SMA exhibited relatively uniform size and morphology, consistent with the corresponding DLS size distributions. The diameters of all PMND structures in each formulation were quantified from the TEM micrographs using ImageJ. The measured diameters across all formulations were within the expected size range for nanodiscs^31,32^ (Figure 2F), approximately 20 nm in diameter. While all four synthesis variants generated PMNDs, their relative yields differed substantially. As a comparative measure of PMND yield, we quantified the number of PMNDs observed per µm^2^ of TEM image. PMNDs derived from SPMVs and SMA exhibited a particle density of approximately 600 PMNDs/µm^2^, compared with approximately 10-60 PMNDs/µm^2^ for the other formulations (Figure 2G).

**Figure 2.**
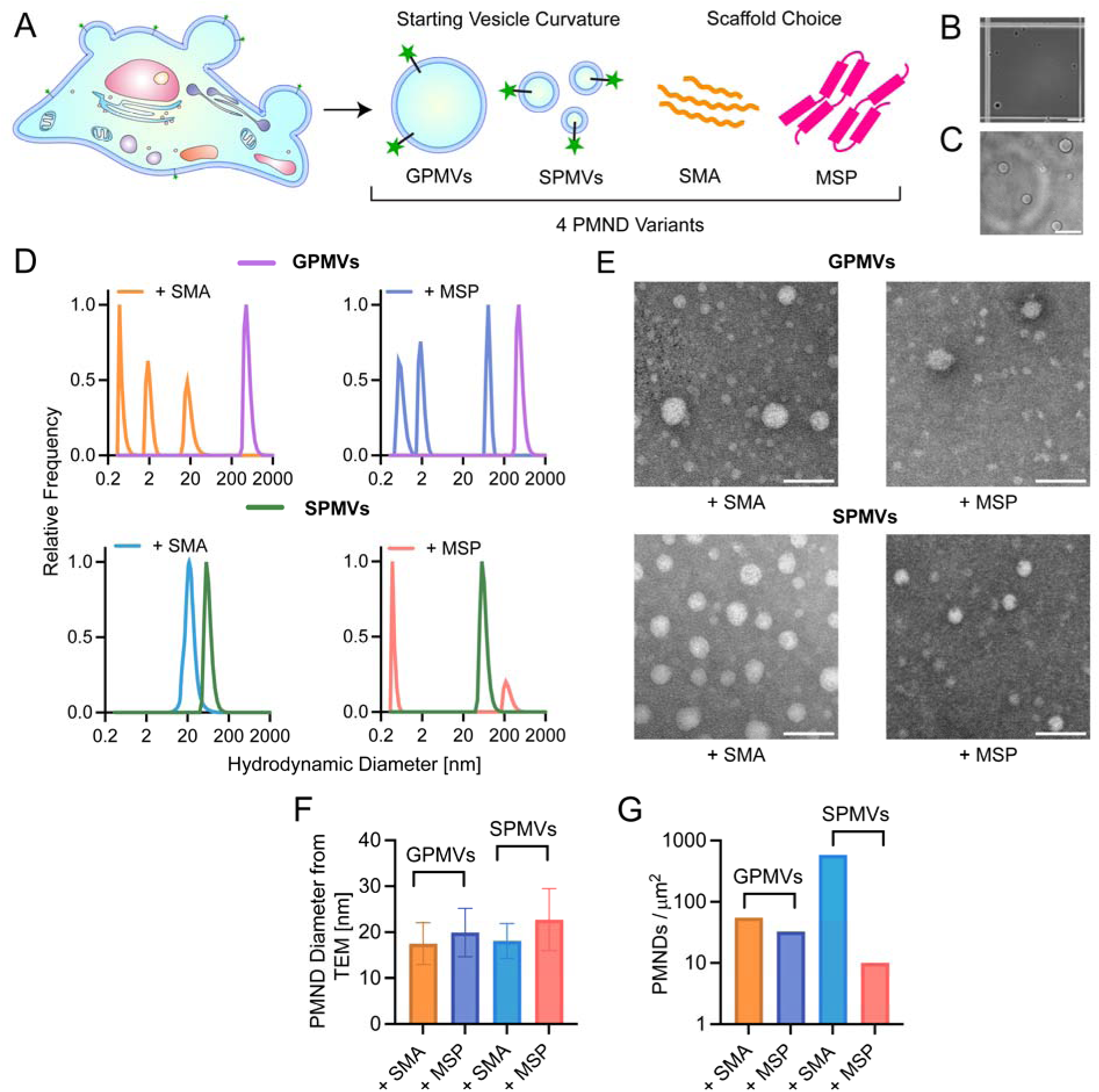
Synthesis optimization for GPMV-derived Plasma Membrane-Enriched Nanodiscs (PMNDs). (A) Schematic of the four starting species for PMND synthesis. Representative brightfield micrographs of (B) purified GPMVs on a hemocytometer grid (scale bar = 40 µm), and (C) purified GPMVs on a glass-bottom dish. (scale bar = 20 µm). (D) DLS size distribution for the four PMND synthesis variants. (E) Representative negative stain TEM micrographs of PMNDs resulting from the synthesis conditions tested (scale bars = 50 nm). (F) Average PMND diameter determined from TEM image analysis. Error bars represent standard error of the mean (SEM). (G) PMND surface density on TEM grids across the four synthesis conditions tested, quantified as the spatial density of nanodisc structures observed on TEM grids.

These results suggest that both scaffold identity and the physical properties of the starting membrane influence PMND formation. MSP-based nanodisc assembly typically relies on detergent-mediated lipid solubilization followed by detergent removal to drive formation of planar bilayer structures stabilized by the scaffold protein.^33^ In contrast, SMA can directly solubilize lipid bilayers through insertion into the hydrophobic membrane core, a process that is enhanced by membrane packing defects.^34^ Because increased membrane curvature promotes lipid splay and packing defects^35^, the smaller, more highly curved membrane structures generated by sonication may facilitate SMA insertion and nanodisc formation. Based on its substantially higher proportion of PMND structures, reduced non-PMND material, and greater relative yield, SPMVs + SMA was selected as the best-performing PMND synthesis condition and used for all subsequent experiments.

We next aimed to evaluate the ability of GPMV starting material to retain functional membrane proteins. To this end, we generated a HeLa cell line stably expressing membrane-anchored green fluorescent protein (GFP) as a proof-of-concept membrane protein. GFP was expressed as a fusion protein linked to a truncation mutant of transferrin receptor (TfR) to ensure plasma membrane display. This construct and other similar ones were previously engineered to anchor fluorescent proteins and receptor-targeting ligands to the plasma membrane for quantitative fluorescence analysis.^36,37^ TfR-GFP cells undergoing active vesiculation were visualized using laser scanning confocal microscopy, and GFP was clearly visible in both the in-tact plasma membrane (Figure 3B left) as well as in budding GPMVs (Figure 3B right). Purified GPMVs isolated from these cells also retained TfR-GFP in their membrane (Figure 3C top row). To ensure that the GFP remained properly folded during the vesiculation and purification process, purified GPMVs containing TfR-GFP were incubated with Alexa Fluor 647-labeled GFP nanobody (AF647-GFPnb), a soluble, single-domain antibody fragment that binds GFP with nanomolar affinity.^38^ Confocal micrographs of TfR-GFP GPMVs incubated with AF647-GFPnb demonstrate clear colocalization of GFP membrane signal with AF647, indicating that GFPnb was associating strongly with the GPMV membrane (Figure 3C bottom row). Control experiments were performed with unmodified GPMVs – derived from wild-type HeLa cells – stained with CellMask Green plasma membrane dye. Unmodified GPMVs exhibited exclusion of AF647-GFPnb as there was no significant overlap in signal between the GPMV membrane and the nanobody-fluorophore complex (Figure 3C middle row). Radial profile scans of each condition were performed, and TfR-GFP GPMVs treated with AF647-GFPnb exhibited overlap in fluorescent signal between both channels (Figure 3D). These results establish that TfR-GFP remains membrane-anchored, fluorescent, and accessible to GFPnb in the solution environment after GPMV isolation.

**Figure 3.**
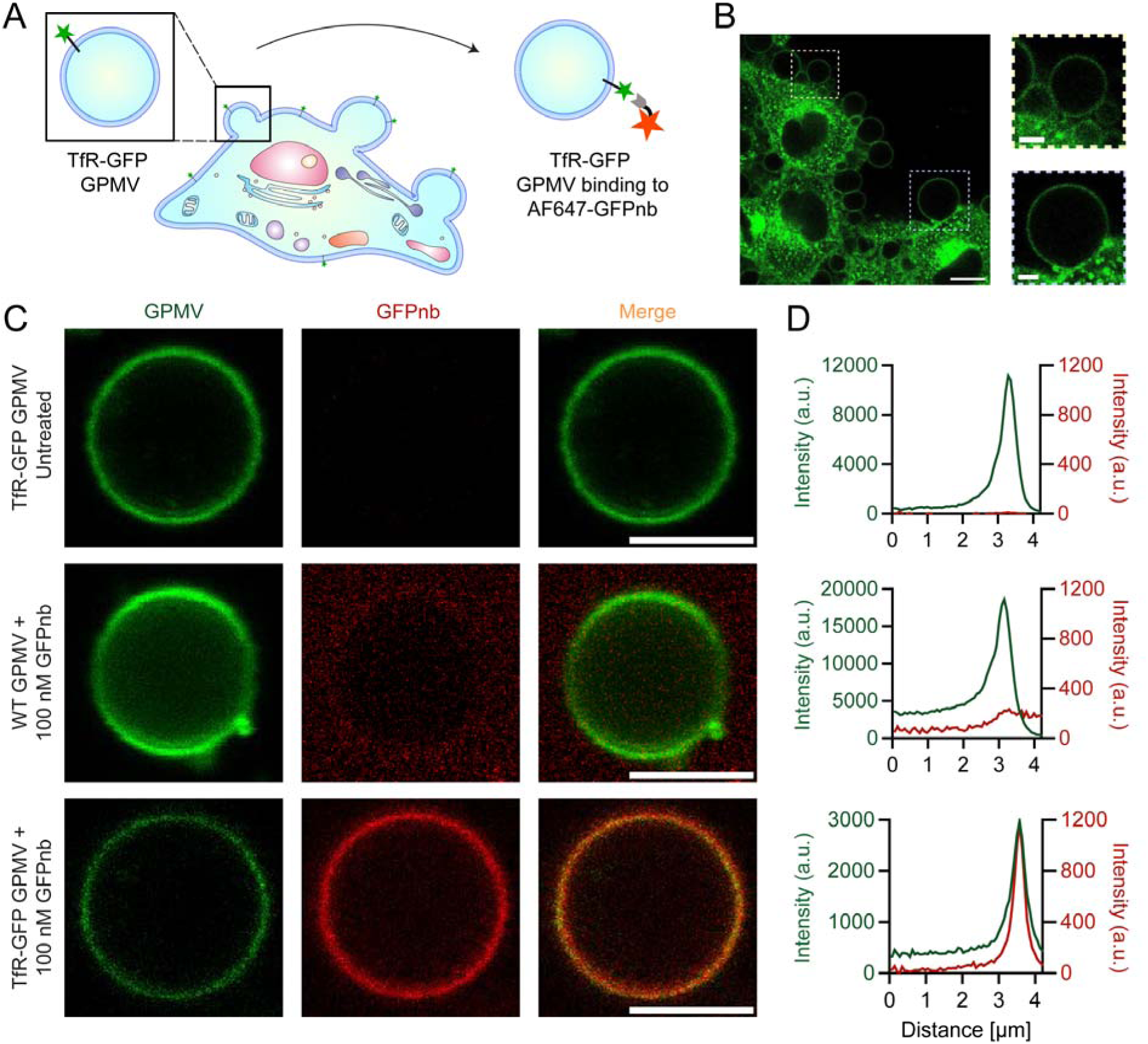
GPMV formation preserves membrane-anchored GFP in a binding-competent state. (A) Schematic of experimental design. (B) Representative confocal micrographs of (left) HeLa TfR-GFP cells undergoing active vesiculation (scale bar = 10 µm) and (right) zoom-ins of active plasma membrane vesiculation (scale bars = 3 µm). (C) Representative confocal micrographs (scale bars = 5 µm) and (D) radial profile scans of untreated TfR-GFP GPMVs (top row), wild-type GPMVs treated with 100 nM AF647-GFPnb (middle row), TfR-GFP GPMVs treated with 100nM AF647-GFPnb (bottom row).

After confirming that GPMVs retained functional TfR-GFP, we synthesized PMNDs from these same TfR-GFP HeLa cells (GFP-PMNDs) and evaluated their interactions with live cell expressing a model receptor containing the GFPnb ligand anchored to the extracellular side of the plasma membrane via TfR and fluorescently labeled with monomeric red fluorescent protein (mRFP) (TfR-mRFP-GFPnb) (Figure 4A). GFP-PMNDs were characterized using DLS, demonstrating a uniform size distribution with an average size of 20 nm (Figure 4B). Using negative stain TEM, GFP-PMNDs were found to exhibit uniform size and morphology (Figure 4C). Then, GFP-PMNDs were incubated with either wild-type HeLa cells or HeLa cells transiently expressing TfR-mRFP-GFPnb. After 20 minutes of incubation, media surrounding the cells was rinsed away and cells were imaged using two-channel confocal microscopy (Figure 4D). Wild-type HeLa cells not expressing the model receptor exhibited negligible GFP fluorescence on the plasma membrane, comparable to the background fluorescence observed in untreated cells expressing the model receptor. In contrast, model receptor-expressing cells treated with 500 nM GFP-PMND exhibited significant GFP fluorescence at the plasma membrane, indicating specific association of the GFP-PMNDs with the model receptor. Line scans of TfR-mRFP-GFPnb expressing HeLa cells treated with 500 nM GFP-PMNDs confirmed strong colocalization of GFP-PMNDs with the mRFP-labeled plasma membrane, which was not observed in wildtype cells (Figures 4E,F). Importantly, GFP-PMND association persisted after removal of excess nanodiscs from the surrounding medium, further supporting stable, receptor-mediated binding rather than transient or nonspecific membrane association.

**Figure 4.**
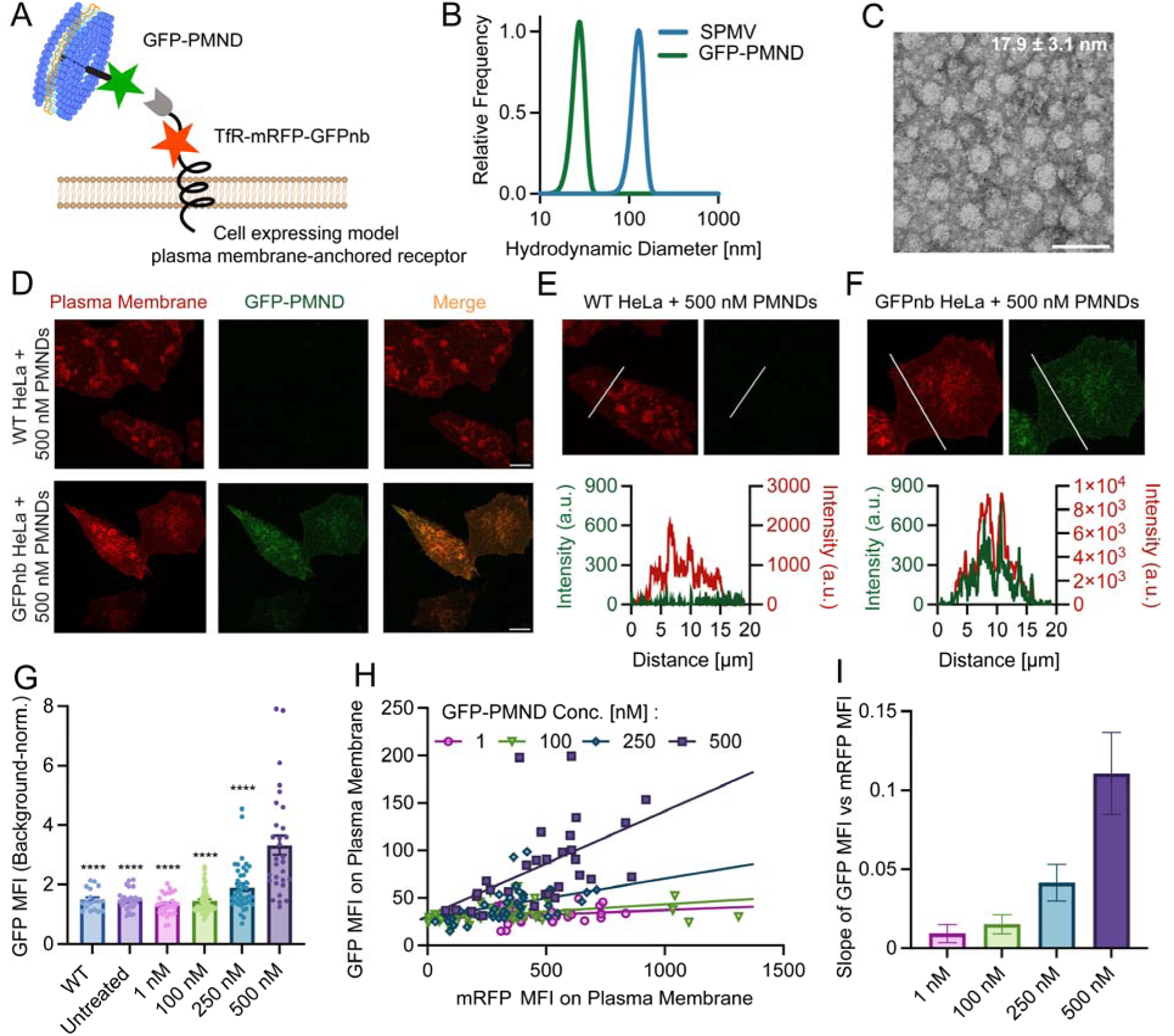
GFP-PMNDs exhibit dose-dependent binding to HeLa cells expressing TfR-mRFP-GFPnb model receptors. (A) Schematic GPMV-derived GFP-PMND binding to cells expressing model plasma membrane-anchored TfR-mRFP-GFPnb receptors. (B) DLS size distributions of SPMVs and GFP-PMNDs. (C) Representative TEM micrograph of GFP-PMNDs (scale bar = 50 nm). Error represents the standard deviation. (D) Representative two-color confocal micrographs of GFP-PMND-treated wild-type HeLa cells (top row) and GFP-PMND-treated HeLa cells expressing TfR-mRFP-GFPnb model receptors (bottom row) (scale bars = 10 µm). Line scans of GFP-PMND-treated (E) wild-type HeLa cells and (F) HeLa cells expressing TfR-mRFP-GFPnb model receptors. (G) GFP-PMNDs exhibit dose-dependent binding to HeLa cells expressing the TfR-mRFP-GFPnb model receptor. WT HeLa cells treated with 500 nM GFP-PMNDs exhibit similar plasma membrane fluorescence to untreated cells expressing the model receptor. Error bars represent SEM. One-way ANOVA shows significant difference between TfR-mRFP-GFPnb-expressing cells treated with 500 nM GFP-PMNDs and all other conditions. ****: p < 0.0001. (H) The mean fluorescence intensity (MFI) of GFP-PMNDs as a function of mRFP MFI on the plasma membrane for various concentrations GFP-PMND incubation concentrations. Solid lines represent linear regressions. (I) The slopes of the linear regressions performed in panel H. Error bars represent 95% confidence intervals.

To quantify this interaction across cells and GFP-PMND concentrations, we next measured the mean fluorescence intensity (MFI) of GFP on the plasma membrane and divided it by the GFP MFI in the background on a per-cell basis, which increased monotonically with GFP-PMND concentration (Figure 4G). Furthermore, GFP-PMND binding increased with model receptor expression levels, as assessed by the relationship between membrane-associated GFP and mRFP MFI (Figure 4H). The slopes of these MFI relationships increased with GFP-PMND concentration from 1 to 500 nM (Figure 4I), indicating progressively greater GFP-PMND binding with increasing receptor abundance and nanodisc concentration. This dependence on both GFP-PMND concentration and receptor abundance is consistent with a receptor-mediated binding process, in which increasing nanodisc concentration promotes receptor occupancy while increasing receptor abundance provides additional binding sites at the cell surface. Taken together, these data demonstrate that TfR-GFP remains fluorescent and accessible to binding partners throughout GPMV isolation and PMND synthesis, enabling specific, sub-micromolar, concentration-dependent binding to cognate receptors on live cells.

## CONCLUSION

In summary, we have successfully developed a method for synthesizing PMNDs derived from GPMVs, providing a streamlined approach for generating nanodiscs from a plasma-membrane enriched starting material. These PMNDs exhibit uniform size and morphology, and they retain membrane-anchored proteins in a functional, binding-competent state for use in cell-based assays. Importantly, probe sonication of the GPMV starting material substantially improved PMND yield, with SMA exhibiting markedly greater compatibility with this detergent-free approach than the protein-based MSP scaffold. These results highlight the importance of both starting membrane curvature and scaffold chemistry in efficient PMND formation. One limitation of the present study is that PMND production relies on GPMVs harvested from adherent cells grown in conventional two-dimensional culture, which constrains the amount of plasma membrane starting material that can be generated at scale. Extending this approach to three-dimensional or other high-density culture formats would substantially increase cell biomass and improve the scalability of PMND production. GPMVs have previously been isolated from cells grown in suspension^39^, so incorporating these approaches could further facilitate larger-scale PMND synthesis. Additionally, future studies should investigate the incorporation and activity of endogenous or physiologically relevant membrane proteins and evaluate the compositional fidelity of the resulting PMNDs relative to the source plasma membrane. Overall, the method presented in this work circumvents many of the shortcomings associated with traditional plasma membrane isolation and enrichment, establishing GPMVs as a versatile starting material for PMND synthesis, with potential applications in membrane protein structural determination, development of novel hybrid biomaterials, and cell-specific targeting.

## MATERIALS AND METHODS

### Materials

Tris(hydroxymethyl)aminomethane (tris), cOmplete Mini EDTA-free protease inhibitor tablets, 2-mercaptoethanol (BME), L-glutathione reduced (GSH), sodium chloride (NaCl), and phenylmethanesulfonylfluoride (PMSF) were purchased from Sigma Aldrich. 4-(2-hydroxyethyl)-1-piparazineethanesulphonic acid (HEPES), Triton X-100, CellMask Deep Red plasma membrane stain, fetal bovine serum (FBS), Alexa Fluor 647 NHS ester dye, GST agarose, Ni-NTA agarose, and imidazole were purchased from Fisher Scientific. Puromycin was purchased from Invitrogen. Isopropyl-beta-D-thiogalactoside (IPTG) and dithiothreitol (DTT) were purchased from Gold Biotechnology. Trypsin (0.05%), penicillin/streptomycin/L-glutamine (PSLG), Dulbecco’s modified Eagle medium (DMEM) high glucose without L-glutamine and sodium pyruvate, and DMEM high glucose without L-glutamine, sodium pyruvate, and phenol red were purchased from Cytiva Life Sciences. FuGENE HD was purchased from Fugent. Dulbecco’s phosphate-buffered saline (DPBS) was purchased from VWR. SMALP200 copolymer (SMA) was purchased from Cube Biotech. Polybrene was obtained from GenScript.

### Plasmids

The pEGFP-N1-TfRΔEctoR1-mRFP-GFPnb plasmid (Addgene #113555) and the pEGFP-N1-TfRΔEctoR1-eGFP A206K (Addgene #113554) were a kind gift from Dr. Jeanne Stachowiak. The pGEX6P1-GFP-Nanobody (GFPnb) plasmid was a kind gift from Dr. Kazuhisa Nakayama (Addgene Plasmid #61838). The pET28a-pMSP1D1 plasmid was a kind gift from Dr. Stephen Sligar (Addgene Plasmid #20061).

All plasmids were propagated in DH5α *E. coli* and purified using QIAprep Spin Miniprep Kit (Qiagen). Plasmids were stored in nuclease-free water (Invitrogen) and sequenced using the Plasmidsaurus standard plasmid sequencing service. All protein expression was carried out in BL21 (DE3) *E. coli*.

### Protein Purification

<u>GFPnb:</u> GFPnb was expressed as a fusion protein containing N-terminal glutathione-S-transferase (GST). BL21 cells were transformed with the pGEX6P1-GFP-Nanobody plasmid, grown in 2xTY pH 7.4, 100 µg mL^-1^ ampicillin, at 37 °C to an OD600 of 0.6, cooled to 30 °C, induced with 1mM IPTG, and expressed for 6 hours. Pellets were frozen and stored at –80 °C. For purification, pellets from 2 L of culture were thawed on ice, suspended in 40 mL of 500 mM Tris, 5 mM EDTA, 5% v/v glycerol, 10 mM BME, 1 mM PMSF, 1% v/v Triton X-100, and 4 tablets of cOmplete Mini EDTA-free protease inhibitor cocktail (Roche), and probe sonicated (Branson Ultrasonics SLPe Digital Sonifier) at 60% amplitude on ice for 4 total 4-minute cycles (2 minutes on, 2 minutes off). The sonicated lysate was then spun down at 103,800 g for 30 minutes at 4 °C. The lysate was then batch absorbed to 2 mL bed volume glutathione agarose resin (Thermo Fisher) for 1 hour at 4 °C with stirring at 60 rpm. The column was then: (1) washed with 10 column volumes (CV) of 400 mM Tris, 1 M NaCl, 10 mM BME, 1 mM PMSF pH 8.0, (2) washed with 10 CV of 25 mM HEPES, 150 mM NaCl, 1 mM EDTA, 10 mM BME pH 7.4 and (3) eluted with 4 CV of 150 mM NaCl, 50 mM Tris, 20 mM GSH, 1 mM EDTA pH 8.0. The eluate was concentrated with a 30 kDa MWCO Amicon Ultra centrifugal concentrator (Millipore) and incubated with soluble HRV-3C protease (Pierce) overnight at 4 °C. Excess HRV-3C, which contained a GST tag, was removed by passage through another glutathione agarose column. The eluate was then concentrated and buffer exchanged into 25 mM HEPES, 150 mM NaCl, 5 mM DTT, pH 7.4 with a 3 kDa MWCO Amicon Ultra centrifugal concentrator (Millipore) operated at 4,000 g. The concentrated protein was then run on a Superdex 75 10/300 size exclusion chromatography column (Cytiva) at 4 °C to remove residual GST. The protein fractions containing GFPnb were pooled, concentrated with a 3 kDa MWCO Amicon Ultra centrifugal concentrator, and incubated with Alexa Fluor 647 NHS ester at a 3:1 dye-to-protein ratio. This mixture reacted at room temperature for 30 minutes, after which a 3 kDa MWCO Amicon Ultra centrifugal concentrator was used to filter the protein and remove excess dye. The final dye:protein ratio was determined to be 1:1 via UV/Vis spectroscopy using a NanoDrop (Thermo Fisher Scientific).

<u>MSP1D1:</u> his-tagged MSP1D1 was expressed and purified as previously described.^32^ Briefly, transformed BL21 cells were grown in 2xTY media, pH 7.4, 100 µg mL^-1^ ampicillin, at 37 °C to an OD600 of 0.6, cooled to 30 °C, induced with 1 mM IPTG, and expressed for 3 hours. Cells were harvested by centrifugation at 9000 g for 20 minutes, resuspended in 500 mM Tris, 5% v/v glycerol, pH 8.0 with cOmplete Protease Inhibitor tablets (Roche), and lysed by probe tip sonication. Clarified lysates were batch adsorbed onto 3 mL (6mL of 50% slurry) of Ni-NTA resin for 1 hour with stirring at 60 rpm before being applied to a gravity-flow column. Unbound proteins were removed by rinsing with 500 mM Tris, pH 8.0 with increasing concentrations of imidazole (0 mM, 10 mM, 20 mM). Protein was eluted with 500 mM Tris, 200 mM imidazole, pH 8.0. All buffers included 10 mM BME, and 20 mM PMSF was added to all buffers except the lysis buffer. BME and PMSF were added to buffers immediately before use. Eluted fractions were pooled and dialyzed for 24 hours against 25 mM HEPES, 150 mM NaCl, pH 7.4 to remove remaining imidazole.

All purified proteins were aliquoted, flash frozen in liquid nitrogen, and stored at –80 °C. Protein molecular weight and purity were determined by SDS-PAGE.

### Cell Culture and Transfection

HeLa cells were a kind gift from Dr. Jeanne Stachowiak (University of Texas, Austin). Cells were cultured in DMEM media supplemented with 10% fetal bovine serum, Penicillin / Streptomycin / L-glutamine (100 units mL^-1^ / 100 µg mL^-1^ / 300 µg mL^-1^ respectively), and incubated at 37 °C with 5% CO_2_.

### Stable Cell Line Generation

To generate a cell line stably expressing membrane-anchored GFP, the TfRΔEctoR1-eGFP A206K plasmid sequence was inserted into a pLVV-CMV-PGK-Puro vector for lentiviral expression. Plasmid construction and lentivirus packaging was completed by GenScript. Upon shipment, the lentivirus was split into single use aliquots in 0.6 mL tubes (Eppendorf).

Purified lentiviruses from GenScript were used to transduce HeLa cells. HeLa cells were seeded at a density of 5 10 cells in a 10 cm dish. After 24 h, the media was replaced with complete growth media containing 8 μg mL^-1^ polybrene. The lentivirus aliquots were then thawed on ice and added to the HeLa cells drop-wise at a multiplicity of infection (MOI) of 30. The cells were incubated with the lentivirus in polybrene-containing media for 24h. After 24h, the cells were rinsed with DPBS, and the media was replaced with regular growth media. After 24h in regular growth media, the media was replaced with media containing 3 μg mL^-1^ puromycin for antibiotic selection. The cells were selected for 4 days, after which they were bright sorted using fluorescence-activated cell sorting.

### Giant Plasma Membrane Vesicle (GPMV) Synthesis

GPMVs were synthesized and purified according to previous protocols, with modifications.^28^ Cells were grown in 175 cm^2^ Nunc EasYFlask Cell Culture Flasks (Thermo Scientific) until they reached 80% confluence. Cells were rinsed twice with 6 mL GPMV Buffer (10 mM HEPES, 150 mM NaCl, 2 mM CaCl_2_, pH 7.4) and once with 6 mL Active GPMV Buffer (GPMV buffer with 2 mM N-Ethylmaleimide added immediately before use). Then, cells were incubated with 6 mL Active GPMV Buffer at 37 °C with 5% CO_2_. After 6-10 hours incubation, GPMVs were carefully collected from flasks by pipette and centrifuged at 700 g for 5 minutes at room temperature to pellet any cell debris. Then, the supernatant was collected and centrifuged at 17,000 g for 20 minutes at 4 °C to pellet GPMVs. The GPMV pellet was resuspended in an appropriate buffer: GPMV Buffer for GPMV experiments or PMND Buffer (25 mM HEPES, 150 mM NaCl, pH 7.4) for PMND experiments.

### Plasma Membrane-Derived Nanodisc (PMND) Synthesis

To generate PMNDs, SMA or MSP was incubated with either GPMVs or SPMVs at a defined scaffold-to-membrane ratio. To determine this ratio, we first approximated the total plasma membrane surface area in each GPMV sample. GPMVs were resuspended in PMND buffer, and the total GPMV surface area was approximated by multiplying the GPMV concentration (i.e., number of GPMVs per unit volume) by the average surface area per GPMV. GPMV concentrations were determined by using a hemocytometer (Hauser Scientific) to count the average number of GPMVs present in a fixed sample volume. Average GPMV diameters were estimated from brightfield microscopy acquired using an ECHO Revolve Microscope and analyzed in ImageJ. Average GPMV surface area was then calculated assuming spherical geometry, such that *area* = *Π* (*diameter*)^2^. SPMVs were generated by performing probe sonication GPMVs at an amplitude of 60% on ice for 1 minute.

We then estimated the minimum number of scaffold molecules required per unit membrane surface area (i.e., the area occupied by a continuous bilayer, not the area each leaflet). Based on a minimum scaffold:nanodisc ratio of 2:1 and an approximate disc diameter of 20 nm, we estimated the minimum scaffold requirement per membrane surface area as follows:

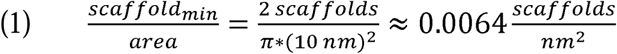

Based on this minimum stoichiometry, we operated at a 30-fold molar excess, incubating SMA or MSP with GPMVs or SPMVs at a ratio of approximately 0.2 scaffold molecules/nm^2^ of membrane surface area. SMA was dissolved in water at 10% w/v prior to incubation with GPMVs or SPMVs. Incubation was carried out overnight at 4 °C with stirring at 60 rpm. The following morning, PMND solutions were ultracentrifuged at 150,000 g for 30 minutes at 4 °C to pellet unreacted material. The PMNDs were further purified and concentrated using a 50 kDa MWCO Amicon Ultra centrifugal concentrators. Size distributions of purified PMNDs were then characterized using dynamic light scattering (DLS) (Horiba SZ-100).

### Transmission Electron Microscopy (TEM)

Samples were prepared on 200 square mesh, formvar-supported, carbon-coated grids (Electron Microscopy Sciences). Grids were cleaned by UV/ozone treatment for 4 minutes. Concentrated PMND solutions were diluted 5-10-fold in PMND buffer. and 5 µL of sample was applied each grid for 1 minute. Excess liquid was wicked away with filter paper (Whatman), and the grids were stained twice with 5 µL of 1% uranyl acetate for 10 seconds per stain. TEM micrographs were collected on a Talos F200C G1 Transmission Electron Microscope using an operating voltage of 80 kV. PMND diameters and particle density were quantified using ImageJ.

### GPMV-GFPnb Binding Experiments

PLL-PEG was synthesized as described previously^37,40^. Imaging wells consisted of 1.6 mm thick silicone gaskets (Grace Bio-Labs) with 5 mm diameter circular holes cut into them. These gaskets were placed directly on top of # 1.5 glass coverslips. Prior to use, gaskets and coverslips were cleaned with Hellmanex (Hellma Analytics) and dried under nitrogen. 30 µL of PLL-PEG was added to wells, allowed to incubate for 20 minutes, and rinsed with GPMV buffer.

Unmodified GPMVs were incubated with CellMask Green plasma membrane stain for 5 minutes, pelleted at 17,000 g for 20 minutes, and resuspended in GPMV buffer to remove excess dye. (1) unmodified GPMVs + 100 nM AF647-GFPnb, (2) TfR-GFP-containing GPMVs + 100 nM AF647-GFPnb, or (3) untreated TfR-GFP-containing GPMVs were incubated in GPMV buffer for 10 minutes; 30 µL of this solution was added to each imaging well, and GPMVs were allowed to settle for 10 minutes before imaging.

### Live Cell Binding Experiments

Imaging wells consisted of 1.6 mm thick silicone gaskets (Grace Bio-Labs) with 5 mm diameter circular holes cut into them. Prior to use, gaskets were cleaned with Hellmanex followed by bath sonication in 70% ethanol. Gaskets were stored in 90% ethanol, and prior to use were dried inside a sterile biosafety cabinet.

Clean, dry gaskets were placed directly on top of 35 mm dishes with a 20 mm #1.5 cover glass micro-well (CellVis). Cells were seeded into imaging wells at a density of 5 10 cells 24 hours prior to transfection. Transfection was carried out using 1 µg of TfR-mRFP-GFPnb plasmid DNA and 3 µL of FuGENE HD reagent, and cells were grown for another 24 hours prior to imaging. Wild-type (WT) cells were seeded 24 hours prior to imaging. Before imaging, WT cells were incubated with 1x CellMask Plasma Membrane Stain Deep Red, which was rinsed with DPBS after 10 minutes.

Nanodiscs were added to imaging wells and incubated at 37 °C for 20 minutes prior to imaging. Cells were washed with DPBS to remove excess nanodisc, media was replaced with phenol red-free DMEM supplemented with 10% FBS, and cells were imaged immediately.

### Laser Scanning Confocal Microscopy

Fluorescence imaging was performed on a laser scanning confocal microscope (Leica Stellaris 5). Three excitation lasers were used: 488 nm for either GFP or CellMask Green plasma membrane stain, 561 nm for mRFP, and 638 nm for either Alexa Fluor 647 or CellMask Deep Red plasma membrane stain. The following emission windows were chosen: 493 – 555 nm for GFP and Cell Mask Green, 576 – 700 nm for mRFP, and 643 – 730 nm for Cell Mask Deep Red and Alexa Fluor 647. A Leica HC PL APO 63x, 1.4 NA oil-immersion objective was used to acquire images, and the zoom factor was set such that 70 nm square pixels were obtained. Cells were imaged at the dorsal plane of the plasma membrane and GPMVs were imaged at the equatorial plane.

### Image Analysis – GPMV Binding

AF647-GFPnb binding to GPMV membranes was quantified using the Radial Profile Plot plugin in ImageJ, which plots normalized integrated intensities around concentric circles as a function of distance. Overlapping intensity distributions in separate channels indicates colocalized fluorescent signal in those channels.

### Image Analysis – Cell Binding

GFP-PMND binding to the cellular plasma membrane was quantified using a custom MATLAB script. Cells were segmented from background signal by defining regions of interest (ROIs) in ImageJ using the mRFP channel, which labels the plasma membrane. Separate masks corresponding to cellular and background regions were generated and applied to the GFP fluorescence channel. The mean fluorescence intensity (MFI) of GFP within each cellular ROI was divided by the mean background GFP fluorescence intensity from the corresponding image to obtain background-normalized GFP fluorescence intensities on a single-cell basis. The mRFP MFI within each cellular ROI was also quantified for the analyses shown in Figures 4H-I. All statistical analyses were performed using GraphPad Prism version 11.

## ACKNOWLEDGEMENTS

Thank you to Dr. Peter Yingxiao Wang’s laboratory at the University of Southern California (USC), especially Dr. Linshan Zhu, for assistance with fluorescence-activated cell sorting. TEM images were obtained at the Core Center for Excellence in Nano Imaging at USC with aid from Carolyn B. Marks. This work was supported in part by the National Institutes of Health through R35GM147333 to C.O.J., B.S.P., and W.F.Z.

## Notes

### Competing Interest Statement

The authors have declared no competing interest.

## REFERENCES

(1) Groves, J. T.; Kuriyan, J. Molecular Mechanisms in Signal Transduction at the Membrane. Nat. Struct. Mol. Biol. 2010, 17 (6), 659–665. 10.1038/nsmb.1844

(2) Hubbard, S. R.; Miller, W. T. Receptor Tyrosine Kinases: Mechanisms of Activation and Signaling. Curr. Opin. Cell Biol. 2007, 19 (2), 117–123. 10.1016/j.ceb.2007.02.010

(3) Guan, L. Structure and Mechanism of Membrane Transporters. Sci. Rep. 2022, 12 (1), 13248, s41598-022-17524–1. 10.1038/s41598-022-17524-1

(4) Jelokhani-Niaraki, M. Membrane Proteins: Structure, Function and Motion. Int. J. Mol. Sci. 2022, 24 (1), 468. 10.3390/ijms24010468

(5) Alberts, B.; Johnson, A.; Lewis, J.; Raff, M.; Roberts, K.; Walter, P. Membrane Proteins. In Molecular Biology of the Cell.

(6) Sligar, S. G.; Denisov, I. G. Nanodiscs: A Toolkit for Membrane Protein Science. Protein Sci. 2021, 30 (2), 297–315. 10.1002/pro.3994

(7) Dalal, V.; Arcario, M. J.; Petroff, J. T.; Tan, B. K.; Dietzen, N. M.; Rau, M. J.; Fitzpatrick, J. A. J.; Brannigan, G.; Cheng, W. W. L. Lipid Nanodisc Scaffold and Size Alter the Structure of a Pentameric Ligand-Gated Ion Channel. Nat. Commun. 2024, 15 (1), 25. 10.1038/s41467-023-44366-w

(8) Pugh, C. F.; Feilen, L. P.; Živković, D.; Præstegaard, K. F.; Sideris, C.; Borthwick, N. J.; De Lichtenberg, C.; Bolla, J. R.; Autzen, A. A. A.; Autzen, H. E. MAASTY: A (Dis)Ordered Copolymer for Structural Determination of Human Membrane Proteins in Native Nanodiscs. Nat. Commun. 2025, 16 (1), 11399. 10.1038/s41467-025-66208-7

(9) Reyes-Ordoñez, A.; Shree, S.; Singh, N.; Sligar, S. G.; Chen, J. Nanodisc Single-Molecule Pulldown to Study Lipid-Protein Interactions. J. Lipid Res. 2025, 66 (7), 100846. 10.1016/j.jlr.2025.100846

(10) Kawai, T.; Caaveiro, J. M. M.; Abe, R.; Katagiri, T.; Tsumoto, K. Catalytic Activity of MsbA Reconstituted in Nanodisc Particles Is Modulated by Remote Interactions with the Bilayer. FEBS Lett. 2011, 585 (22), 3533–3537. 10.1016/j.febslet.2011.10.015

(11) Bayburt, T. H.; Vishnivetskiy, S. A.; McLean, M. A.; Morizumi, T.; Huang, C.; Tesmer, J. J. G.; Ernst, O. P.; Sligar, S. G.; Gurevich, V. V. Monomeric Rhodopsin Is Sufficient for Normal Rhodopsin Kinase (GRK1) Phosphorylation and Arrestin-1 Binding. J. Biol. Chem. 2011, 286 (2), 1420–1428. 10.1074/jbc.M110.151043

(12) Noh, I.; Guo, Z.; Wang, R.; Zhu, A. T.; Krishnan, N.; Mohapatra, A.; Gao, W.; Fang, R. H.; Zhang, L. Modular Functionalization of Cellular Nanodiscs Enables Targeted Delivery of Chemotherapeutics into Tumors. J. Controlled Release 2025, 378, 145–152. 10.1016/j.jconrel.2024.12.004

(13) Sarcinella, M. C.; Jones, J. D.; Sorensen, M. J.; Edgcombe, S. A.; Ruotolo, B. T.; Kennedy, R. T.; Bailey, R. C. Lipid Curvature and Fluidity Influence Lipid Incorporation Disparities in Nanodiscs. Anal. Chem. 2025, 97 (5), 2883–2889. 10.1021/acs.analchem.4c05494

(14) Knetsch, T. G. J.; Van Son, H.; Kobayashi, M.; Ubbink, M. The Nature of Nanodisc Lipids Influences Fragment Based Drug Discovery Results. Chem. Biol. Drug Des. 2025, 105 (3), e70080. 10.1111/cbdd.70080

(15) Denisov, I. G.; Grinkova, Y. V.; Lazarides, A. A.; Sligar, S. G. Directed Self-Assembly of Monodisperse Phospholipid Bilayer Nanodiscs with Controlled Size. J. Am. Chem. Soc. 2004, 126 (11), 3477–3487. 10.1021/ja0393574

(16) Brown, C.; Ghosh, S.; McAllister, R.; Kumar, M.; Walker, G.; Sun, E.; Aman, T.; Panda, A.; Kumar, S.; Li, W.; Coleman, J.; Liu, Y.; Rothman, J. E.; Bhattacharyya, M.; Gupta, K. A Proteome-Wide Quantitative Platform for Nanoscale Spatially Resolved Extraction of Membrane Proteins into Native Nanodiscs. Nat. Methods 2025, 22 (2), 412–421. 10.1038/s41592-024-02517-x

(17) Guo, Z.; Noh, I.; Zhu, A. T.; Yu, Y.; Gao, W.; Fang, R. H.; Zhang, L. Cancer Cell Membrane Nanodiscs for Antitumor Vaccination. Nano Lett. 2023, 23 (17), 7941–7949. 10.1021/acs.nanolett.3c01775

(18) Sun, L.; Yu, Y.; Peng, Y.; Wang, D.; Wang, S.; Noh, I.; Fang, R. H.; Gao, W.; Zhang, L. Platelet Membrane Derived Nanodiscs for Neutralization of Endogenous Autoantibodies and Exogenous Virulence Factors. Small 2024, 20 (18), 2308327. 10.1002/smll.202308327

(19) Knowles, T. J.; Finka, R.; Smith, C.; Lin, Y.-P.; Dafforn, T.; Overduin, M. Membrane Proteins Solubilized Intact in Lipid Containing Nanoparticles Bounded by Styrene Maleic Acid Copolymer. J. Am. Chem. Soc. 2009, 131 (22), 7484–7485. 10.1021/ja810046q

(20) Sun, L.; Feng, K.; Bai, D.; Yu, Y.; Shen, W.-T.; Zhang, J. A.; Fang, R. H.; Gao, W.; Zhang, L. Hepatic Endoplasmic Reticulum-Derived Nanodiscs for Broad-Spectrum Drug Detoxification. Biomaterials 2025, 318, 123188. 10.1016/j.biomaterials.2025.123188

(21) Goldberg, S. Mechanical/Physical Methods of Cell Disruption and Tissue Homogenization. In *Proteomic Profiling*; Posch, A., Ed.; Methods in Molecular Biology; Springer US: New York, NY, 2021; Vol. 2261, pp 563–585. 10.1007/978-1-0716-1186-9_36

(22) Sandelius, A. S.; Morré, D. J. Plasma Membrane Isolation. In The Plant Plasma Membrane; Larsson, C., Møller, I. M., Eds.; Springer Berlin Heidelberg: Berlin, Heidelberg, 1990; pp 44–75. 10.1007/978-3-642-74522-5_3

(23) Schimmel, S. D.; Kent, C.; Bischoff, R.; Vagelos, P. R. Plasma Membranes from Cultured Muscle Cells: Isolation Procedure and Separation of Putative Plasma-Membrane Marker Enzymes. Proc. Natl. Acad. Sci. 1973, 70 (11), 3195–3199. 10.1073/pnas.70.11.3195

(24) DePierre, J. W.; Karnovsky, M. L. PLASMA MEMBRANES OF MAMMALIAN CELLS. J. Cell Biol. 1973, 56 (2), 275–303. 10.1083/jcb.56.2.275

(25) De Weerd, S.; Ruiter, E. A.; Calicchia, E.; Portale, G.; Schuringa, J. J.; Roos, W. H.; Salvati, A. Optimization of Cell Membrane Purification for the Preparation and Characterization of Cell Membrane Liposomes. Small Methods 2024, 8 (12), 2400498. 10.1002/smtd.202400498

(26) Boone, C. W.; Ford, L. E.; Bond, H. E.; Stuart, D. C.; Lorenz, D. ISOLATION OF PLASMA MEMBRANE FRAGMENTS FROM HELA CELLS. J. Cell Biol. 1969, 41 (2), 378–392. 10.1083/jcb.41.2.378

(27) Hamilton, A. K.; Mooney, B.; Glisovic-Aplenc, T.; Radaoui, A. B.; Conkrite, K. L.; De Jong, C.; Sidoli, S.; Sorensen, P. H.; Garcia, B. A.; Aplenc, R.; Maris, J. M.; Morin, G. B.; Diskin, S. J. Protocol for Plasma Membrane Enrichment by Ultracentrifugation for Mass Spectrometry of Cell Lines, Xenografts, and Patient Tumors. STAR Protoc. 2025, 6 (3), 104049. 10.1016/j.xpro.2025.104049

(28) Sezgin, E.; Kaiser, H.-J.; Baumgart, T.; Schwille, P.; Simons, K.; Levental, I. Elucidating Membrane Structure and Protein Behavior Using Giant Plasma Membrane Vesicles. Nat. Protoc. 2012, 7 (6), 1042–1051. 10.1038/nprot.2012.059

(29) Scott, R. E. Plasma Membrane Vesiculation: A New Technique for Isolation of Plasma Membranes. Science 1976, 194 (4266), 743–745. 10.1126/science.982044

(30) Baumgart, T.; Hammond, A. T.; Sengupta, P.; Hess, S. T.; Holowka, D. A.; Baird, B. A.; Webb, W. W. Large-Scale Fluid/Fluid Phase Separation of Proteins and Lipids in Giant Plasma Membrane Vesicles. Proc. Natl. Acad. Sci. 2007, 104 (9), 3165–3170. 10.1073/pnas.0611357104

(31) Craig, A. F.; Clark, E. E.; Sahu, I. D.; Zhang, R.; Frantz, N. D.; Al-Abdul-Wahid, M. S.; Dabney-Smith, C.; Konkolewicz, D.; Lorigan, G. A. Tuning the Size of Styrene-Maleic Acid Copolymer-Lipid Nanoparticles (SMALPs) Using RAFT Polymerization for Biophysical Studies. Biochim. Biophys. Acta BBA –Biomembr. 2016, 1858 (11), 2931–2939. 10.1016/j.bbamem.2016.08.004

(32) Pizarro Carbajal, B. S.; Reinhardt, T. G.; Semenske, J. A.; Ji, Z.; Jacobs, C. O.; Zeno, W. Amphiphilic Peptide Fusion Promotes Endocytic Uptake of Nanodiscs. ACS Appl. Bio Mater. 2026, 9 (20), 9656–9669. 10.1021/acsabm.6c01154

(33) Camp, T.; Sligar, S. G. Nanodisc Self-Assembly Is Thermodynamically Reversible and Controllable. Soft Matter 2020, 16 (24), 5615–5623. 10.1039/D0SM00336K

(34) Scheidelaar, S.; Koorengevel, M. C.; Pardo, J. D.; Meeldijk, J. D.; Breukink, E.; Killian, J. A. Molecular Model for the Solubilization of Membranes into Nanodisks by Styrene Maleic Acid Copolymers. Biophys. J. 2015, 108 (2), 279–290. 10.1016/j.bpj.2014.11.3464

(35) Cui, H.; Lyman, E.; Voth, G. A. Mechanism of Membrane Curvature Sensing by Amphipathic Helix Containing Proteins. Biophys. J. 2011, 100 (5), 1271–1279. 10.1016/j.bpj.2011.01.036

(36) DeGroot, A. C. M.; Busch, D. J.; Hayden, C. C.; Mihelic, S. A.; Alpar, A. T.; Behar, M.; Stachowiak, J. C. Entropic Control of Receptor Recycling Using Engineered Ligands. Biophys. J. 2018, 114 (6), 1377–1388. 10.1016/j.bpj.2018.01.036

(37) Zeno, W. F.; Baul, U.; Snead, W. T.; DeGroot, A. C. M.; Wang, L.; Lafer, E. M.; Thirumalai, D.; Stachowiak, J. C. Synergy between Intrinsically Disordered Domains and Structured Proteins Amplifies Membrane Curvature Sensing. Nat. Commun. 2018, 9 (1), 4152. 10.1038/s41467-018-06532-3

(38) Kubala, M. H.; Kovtun, O.; Alexandrov, K.; Collins, B. M. Structural and Thermodynamic Analysis of the GFP:GFP nanobody Complex. Protein Sci. 2010, 19 (12), 2389–2401. 10.1002/pro.519

(39) Tao, X.; Zhao, C.; MacKinnon, R. Membrane Protein Isolation and Structure Determination in Cell-Derived Membrane Vesicles. Proc. Natl. Acad. Sci. 2023, 120 (18), e2302325120. 10.1073/pnas.2302325120

(40) Zeno, W. F.; Thatte, A. S.; Wang, L.; Snead, W. T.; Lafer, E. M.; Stachowiak, J. C. Molecular Mechanisms of Membrane Curvature Sensing by a Disordered Protein. J. Am. Chem. Soc. 2019, 141 (26), 10361–10371. 10.1021/jacs.9b03927

